# Tracking demographic changes in individual coral colonies through repeated photogrammetric surveys

**DOI:** 10.64898/2026.09.29.755387

**Authors:** Kai L. Kopecky, Gaia Pavoni, Massimiliano Corsini, Erica Nocerino, Fabio Menna, Andrew J. Brooks, Russell J. Schmitt, Sally J. Holbrook

**Author notes:** Corresponding author: Kai L. Kopecky.

## Abstract

Understanding the consequences of global change for Earth’s ecosystems requires reliable, accurate, and scalable tools for detecting ecological change. To this end, recent efforts have focused on scaling up the quantification of demographic processes from individual organisms (the fundamental ecological unit) to the larger spatial scales they inhabit to yield more precise and informative estimations of ecological change, especially in response to disturbance. Here, we describe a transferable workflow that combines robust underwater photogrammetry techniques with machine-learning-assisted image analysis to quantify demographic changes in coral colonies—growth, stasis, partial mortality, and complete mortality—through time on the reef landscape. We then demonstrate several analytical applications of this workflow that can be broadly applied to sessile and/or clonal organisms in other ecosystems. First, we tracked the fates of 3,162 coral colonies through a marine heatwave that caused substantial mortality. Second, our approach enabled detection of size- and taxon-dependent patterns in demographic changes from before to after the heatwave. Finally, for a subset (425) of these colonies, we further demonstrated how our method can be used to track fates across multiple years, including a year of change before and after the heatwave took place, revealing nuanced patterns of survival and mortality, as well as changes in size structure that occurred before and after the disturbance. Our workflow offers an important advance for increasing the accuracy of change detection on coral reefs and other ecosystems by linking demographic changes in individual organisms with spatially extensive image-based surveys through time.

## INTRODUCTION

Global change is reshaping Earth’s ecosystems at unprecedented rates and scales. Environmental disturbances are changing in intensity, frequency, and type (Dai, 2013; Gaiser et al., 2020; Johnstone et al., 2016; Oliver et al., 2018), precipitating large and unknown impacts to the structure and function of future ecosystems. Efficient and reliable tools are needed to accurately quantify ecosystem responses to disturbances and help predict the longevity of associated impacts therein. Technologies associated with remote sensing offer much promise in this regard, as they have greatly increased the spatial scales and resolutions at which observations of the environment can be made (Kerr & Ostrovsky, 2003). Bottlenecks associated with extracting data from remote sensing products can potentially be reduced by automating labor- and time-intensive tasks via machine learning (ML), further enhancing the capacity for investigating ecological change (Janga et al., 2023; Zhang & Zhang, 2022).

At the intersection of remote sensing, ML, and ecology is an emerging effort to scale up the quantification of demographic processes (i.e., rates of growth, mortality/survival, and colonization/recruitment) from individual organisms to populations at landscape scales. For example, recent studies in terrestrial forests have measured characteristics and traits of individual trees within airborne remote sensing imagery, then implemented ML models to apply these measurements to individual trees across larger tracts of forest (Marconi et al., 2021) and over multiple years (Battison et al., 2024). This coupling of remote sensing with ML approaches provides an avenue to efficiently link measures of demographic processes in individuals (the fundamental ecological unit) with the broader spatial areas they occupy, increasing the resolution of ecological change detection without compromising spatial scale.

Similar approaches are now emerging in shallow subtidal ecosystems, particularly for ecologically important space holders (Rowan & Kalacska, 2021). For example, drone- and satellite-based imagery combined with ML methods are being used to identify and track salt marsh plants and clonal seagrass patches, enabling landscape-scale estimates of plant stress, density and spatial turnover through time (Meister & Qu, 2024; Ventura et al., 2023). The use of remote sensing to quantify ecological change in underwater environments is expanding particularly rapidly in coral reef ecosystems (Lyons et al., 2020; Remmers, Boutros, et al., 2024; Remmers, Grech, et al., 2024; Stone et al., 2025). Not only are coral reefs one of the most diverse ecosystems on the planet (Knowlton et al., 2010), they are also among the most threatened (Souter et al., 2020). As a result, coral reefs are undergoing drastic alterations in structure and function on a global scale (Hughes et al., 2017). These changes warrant an urgent need for tools and methods that enable us to reliably quantify these transitions and better predict potential ecological consequences.

Underwater photogrammetry—the use of photographs to extract measurements—is a remote sensing tool that has been increasingly implemented to quantify the abundance and cover of benthic organisms in coral reef ecosystems. Typically, this tool has been used to measure temporal changes in coral cover among patches tens to hundreds of square meters in size (Burns et al., 2015; Nocerino et al., 2020; Rossi et al., 2020; Urbina-Barreto et al., 2021) or at the resolution of individual colonies within much smaller areas (Curtis et al., 2023; Ferrari et al., 2017; Lange & Perry, 2020; Morais et al., 2022). Recent advances, however, are increasingly bridging these scales. Large-area imaging has been used to map the size and spatial distribution of thousands of individual coral colonies (Pedersen et al., 2019), while fine-scale repeated photogrammetry can resolve recruitment and succession of benthic organisms through time (Gouezo et al., 2023). More recently, ML approaches have automated the segmentation and classification of benthic organisms within large-area photogrammetric surveys (Remmers et al., 2025), and photogrammetry has enabled measurements of colony size and bleaching severity for thousands of colonies across spatially extensive reef surveys (Álvarez-Noriega et al., 2025).

Together, these advances demonstrate the growing capacity of photogrammetry to resolve individual organisms across increasingly large spatial extents. An important next step is to link this capacity with repeated surveys that preserve the identities of individual colonies through time, enabling demographic fates—such as growth and mortality—to be quantified across spatially extensive reef areas.

Tracking demographic processes in coral colonies has an added complexity relative to typical non-colonial organisms. While a coral colony can grow, remain relatively stable in size (stasis), or die completely from one period to the next, it also can undergo ‘partial mortality’, wherein some living tissue dies and leaves behind dead skeleton along with a surviving portion of the colony (Jackson & Hughes, 1985). Thus, demographic responses of coral colonies may not always be adequately characterized as a binary outcomes of either survival or mortality, but may include varying degrees of tissue loss among surviving colonies. While partial mortality has long been recognized and quantified in the study of coral reefs, effectively quantifying this type of demographic change for large numbers of colonies poses a challenge. Modern image analysis techniques like *semantic image segmentation*—the delineation and classification of objects within images—enable the parsing of live and dead coral tissue within colonies, providing a tool for accurately measuring partial mortality. When done manually, however, image segmentation is a meticulous and time-consuming process, rendering it infeasible for extensive reef areas captured within large photomosaics that can contain hundreds or even thousands of coral colonies (C. Burns et al., 2022; Kopecky, Pavoni, et al., 2023; Pavoni et al., 2022; Sauder et al., 2024; Schürholz & Chennu, 2023). Using deep ML approaches to automate image segmentation can dramatically reduce the time and labor required for this task (Kopecky, Pavoni, et al., 2023; Pavoni et al., 2022), substantially expediting the throughput of image analysis and quantification of nuanced demographic changes for large numbers of coral colonies.

Here, we describe a workflow that integrates photogrammetric surveys, ML-assisted segmentation, and temporally repeated mapping of coral reefs to track demographic changes in individual organisms across repeated reef surveys. We demonstrate how our workflow can be used to measure, identify, and track the fates of thousands of coral colonies, both in response to a major disturbance—a marine heatwave that caused widespread coral bleaching and elevated mortality of coral tissue—as well as in a non-disturbance period. Our previous work described how this image-processing workflow improves estimations of disturbance-driven losses in coral cover by measuring changes in three dimensions as opposed to two (Kopecky, Pavoni, et al., 2023). We build further on this here by preserving colony identities across repeated surveys, allowing individual-level demographic changes to be quantified across more spatially extensive surveys. Specifically, we illustrate how our workflow can be used to quantify colony-level growth, stasis, and mortality (both partial and complete). We also demonstrate the capacity for our workflow to detect size- and taxon-dependent patterns, as well as changes in the size structure and taxonomic composition of coral communities. Importantly, the techniques we describe can also be applied to other sessile and/or clonal organisms in aquatic or terrestrial ecosystems, especially those subject to partial mortality or size shrinkage. Our workflow provides a means to improve monitoring efforts on coral reefs, advance studies of coral reef demography, more accurately evaluate management actions, and better predict downstream consequences of disturbance in benthic communities.

## MATERIALS AND METHODS

### Study site

Moorea, French Polynesia (17°30′S, 149°50′W) is a high volcanic island in the South Pacific Ocean with a barrier reef surrounding the island’s ∼60 km perimeter, from which steep fore reef slopes extend offshore. The Moorea Coral Reef Long Term Ecological Research (MCR LTER, https://mcr.lternet.edu) program has been collecting time series data on coral reef communities around the island since 2005. More recently, photogrammetric survey methods have been developed and implemented to enable close tracking of benthic dynamics, particularly in response to disturbance events such as cyclones and coral bleaching (Capra et al., 2017; Guo et al., 2016; Kopecky, Pavoni, et al., 2023; Neyer et al., 2018; Nocerino et al., 2020; Rossi et al., 2020). In April 2019, a marine heatwave caused prolonged elevated sea surface temperatures that resulted in a major coral bleaching event, ultimately causing substantial mortality of *Pocillopora* and *Acropora* on the fore reef (Burgess et al., 2021; Speare et al., 2022). Our photogrammetric surveys spanned this major disturbance, providing a baseline before the event (2017-2018) and capturing the subsequent, delayed impacts after heat stress had subsided (2019-2020). Below, we outline our method and workflow step by step (Fig. 1).

**Figure 1.**
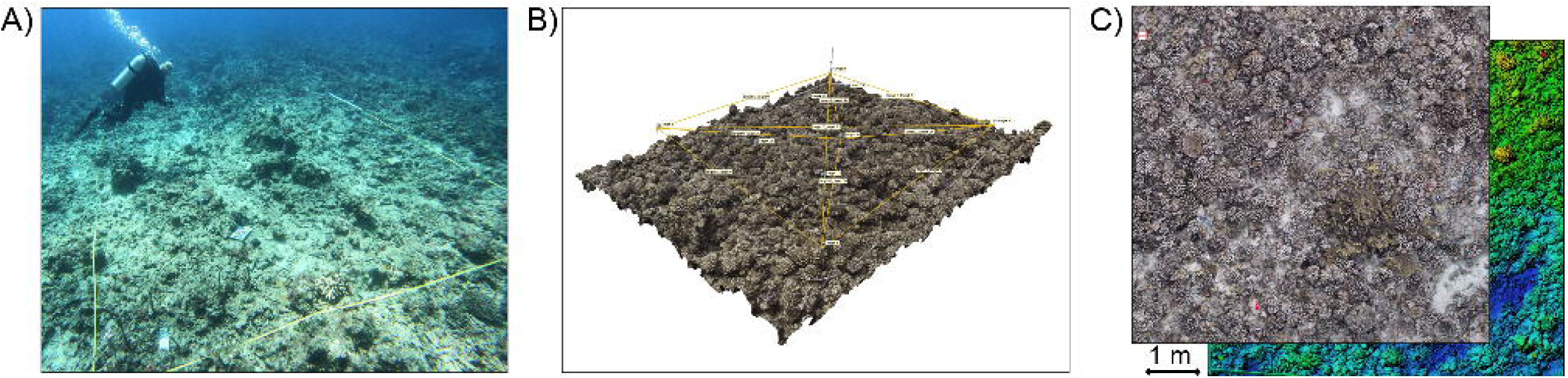
Diagram of the photogrammetric portion of our workflow. (A) A diver collecting photographs of a long-term reef monitoring plot. (B) A 3-D model of a long-term reef monitoring plot with the geodetic network (distances between permanent reference points) displayed. (C) An orthophoto mosaic of one of our reef plots overlaid on a Digital Elevation Model (DEM) of the same plot.

### Establishment of a geodetic network and photogrammetric surveys

Beginning in 2017, we used a robust method of underwater photogrammetry developed by Nocerino et al. (2020) to annually survey five ∼25 m^2^ reef plots (Fig. 1A). By utilizing a ‘geodetic network’ of fixed reference points installed in the reef matrix, this technique enables millimeter-scale change detection of objects within the bounds of the network and precise spatial co-registration of Digital Elevation Models (DEMs) and orthorectified photomosaics (hereafter, *orthophoto mosaics*) through time. To establish this network, we installed permanent threaded anchors (affixed with marine epoxy) into the corners of each 25 m^2^ plot, as well as several additional reference points inside the perimeter set by these corners. Distance and height difference measurements were then taken between all fixed reference points, achieving millimetric estimated accuracy (see Capra et al., 2017; Nocerino et al., 2020).

When conducting our photogrammetric surveys of each plot, we first screwed coded photogrammetry targets (obtained from AgiSoft Metashape, version 2.0) adhered atop metal posts into the threaded, permanent anchors. The posts help elevate the coded targets above the coral canopy. This is needed to ensure visibility between targets and allow for measurement of distances between targets using measuring tapes. The same spatial orientation of targets and posts were used at each sampling, which supports the image orientation step in Metashape and allows for setting reference of the 3D models in a local coordinate system (i.e., the geodetic network; Fig. 1B). SCUBA divers then swam fixed ‘lawnmower’ patterns (a series of parallel passes followed by another series of passes perpendicular to the first), continuously taking photographs ∼1 m above the reef. Divers collected oblique images by swimming around the circumference of each plot and making several passes within each plot. This process ensured sub-millimeter-scale ground sample distance (pixel size expressed in ground/object space units). Roughly 1000-1500 photographs were taken of each of the five 25 m^2^ reef plots in each year from 2017 to 2020.

### Construction of DEMs and orthophoto mosaics

Using Metashape, we constructed 3-D models of each of the five 25 m^2^ reef plots in each year (Fig. 1B). The coordinates of the geodetic network were assumed fixed and stable over time, providing a common metric coordinate reference system to align models across time points and enable the detection and monitoring of temporal changes in reef organisms via photogrammetry. We then used our 3-D models to construct DEMs and orthophoto mosaics of each plot (Fig. 1C) in each year for use in the image segmentation process steps described below, exporting each at a set resolution of 0.5mm/pixel.

### Image segmentation and annotation of coral colonies using TagLab

We used the open-source, automated image segmentation software TagLab (Pavoni et al., 2022) to segment and identify (i.e., annotate) live and dead coral colonies within the orthophoto mosaics (Fig. 2A). TagLab includes a supervised learning workflow consisting of the creation of a training dataset, the training of a semantic segmentation network, and the testing of predictions on new orthophoto mosaics. We first built a training dataset from two of the orthophoto mosaics--one from before the 2019 heatwave and one following it—to capture variation in the sizes and amounts of live and dead coral. The training dataset was built using TagLab’s AI-based interactive tracing tools, which offer rapid point-and-click methods for annotating coral colonies. From this training dataset, we fine-tuned a fully automatic semantic segmentation model to annotate live *Pocillopora* colonies and dead branching coral (Kopecky, Pavoni, et al., 2023).

**Figure 2.**
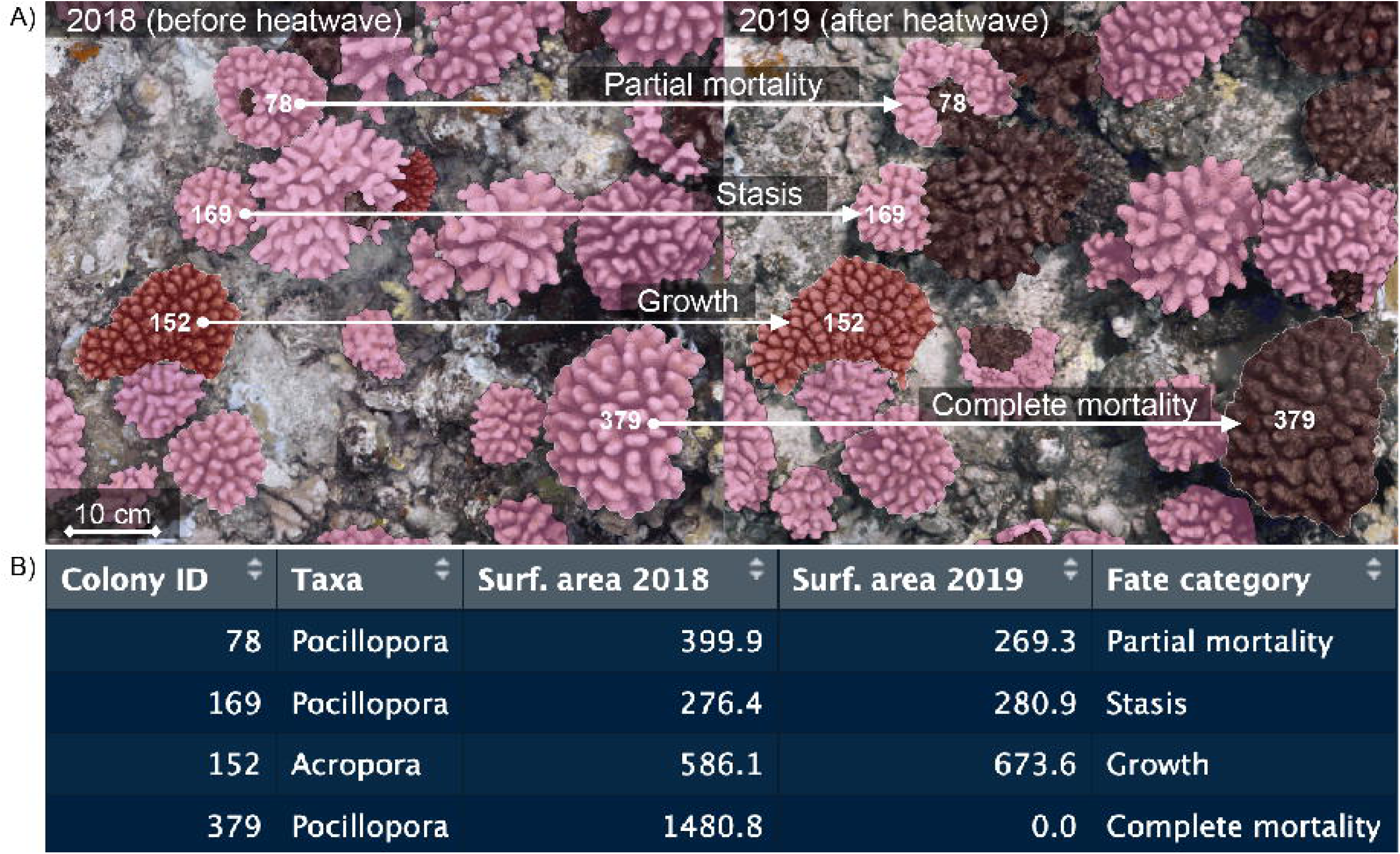
Example of the ML-assisted image analysis workflow in the TagLab interface. (A) A subset of colonies that underwent the four distinct demographic fates (marked by white arrows) from 2018 (before the heatwave) to 2019 (after the heatwave). (B) A data table displaying output from the automated classification of the four colonies outlined in (A), including each colony’s genus, 3-D surface area, unique Colony ID (preserved across time points), and associated demographic fate (Fate Category) over the pre- to post-disturbance period.

Finally, we used our fine-tuned model to automatically segment the remaining orthophoto mosaics. The semantic segmentation network achieved an accuracy of 92% for *Pocillopora* pixels and 71% for dead branching coral pixels, relative to a human observer. The lower accuracy for the dead coral class is primarily due to limitations of the training dataset: only one of the two orthophoto mosaics used for training (the one surveyed after the heatwave) contained a substantial number of dead coral colonies. Moreover, dead corals typically exhibit less sharply defined boundaries and can resemble background substrate, leading to increased classification uncertainty. As a result, 21.9% of dead coral pixels were misclassified as background. TagLab’s graphical interface and post-processing editing tools allowed us to rapidly correct classification errors (< one hour of manual corrections per orthophoto mosaic), thereby improving the quality and accuracy of the final annotated dataset used for subsequent statistical analyses. Because the automatic classification algorithm and its performance evaluation were previously described in detail, we do not repeat those validation analyses here; see Kopecky, Pavoni, et al. (2023) for details on model training, classification accuracy, and manual correction. Lastly, we note that colonies of *Acropora* spp. were relatively rare in our plots and thus were too sparse to provide sufficient training data for a fully automatic classifier, so we annotated *Acropora* colonies using the semi-automatic segmentation tools.

We focused our analyses across the five reef plots at two time points: August 2018, about 8 months before the massive heatwave event when live coral cover was at a maximum, and August 2019, approximately 4 months after the heatwave once significant coral mortality had occurred. We then explored an extended sequence of orthophoto mosaics from 2017 to 2020 for a single reef plot to demonstrate further uses of our method (details provided in the following section).

### Tracking coral colony fates via spatial co-registration through time

Our photogrammetric workflow allows for accurate spatial co-registration of orthophoto mosaics and DEMs of the same reef plot over time (Kopecky, Pavoni, et al., 2023; Nocerino et al., 2020). TagLab exploits this spatial co-registration to assign permanent identifiers to objects that maintain the same class (i.e., live *Pocillopora* or *Acropora*) and physical position through successive time points. In this study, leveraging the open-source nature of TagLab, we implemented a modified version of the temporal tracking algorithm that relaxes the taxonomic class constraint, allowing live and dead specimens of the same species to be tracked. For example, living *Pocillopora* colonies can be matched with dead *Pocillopora* colonies that share the same spatial positions. These matches enable temporal changes to be assigned to one of four predefined categories (hereafter referred to as “fates”; Fig. 2B): *‘Growth’* indicates colonies that increased in size over time with no signs of mortality; *‘Complete mortality’* refers to colonies that had a surface area of live tissue of zero in a later time point (colonies that were present at an earlier time point but no longer detectable at the subsequent time point were classified as such, regardless of whether loss resulted from tissue mortality or physical dislodgement); *‘Partial mortality’* refers to colonies that lost some but not all living tissue over two successive time points; and *‘Stasis’* indicates colonies that changed in size by an amount that fell within our margin of error (∼ 3mm; Nocerino et al., 2020) and therefore could not be reliably classified as having grown or shrunk. TagLab automatically assigns a fifth category for objects that were not detected in a previous time point and appeared in a later one. We omitted 8 instances assigned to this category from our analyses altogether, as it does not always represent newly recruited colonies, but sometimes objects that were simply not detected in the previous time point (e.g., a displaced coral branch that broke off from a larger colony) and therefore require further visual assessment. Colonies < 20 cm^2^ in surface area were omitted from the analyses, as it was difficult even for a human observer to determine the class of objects smaller than this size with the resolution of our orthophoto mosaics. Automated temporal matches and assigned fates were visually inspected against the co-registered orthophoto mosaics and manually corrected where necessary.

### Example analyses of coral mortality and reef change enabled by our method

#### Tracking colony fates through a disturbance event

Across our five replicate reef plots (totaling ∼125 m^2^ in area), we tracked the fates of 3,162 colonies of *Pocillopora* and *Acropora* during the period over which the heatwave occurred (2018-2019). We quantified the number that underwent complete or partial mortality, were static, or grew. We also evaluated the total changes in surface area of live coral associated with each of these categories. Surface area here refers to the colony surface area computed from the DEM, which accounts for only that visible from the top-down perspective of our orthophoto mosaics, rather than the total coral surface area that includes under-parts of the colony that are not visible from this perspective.

#### Influences of colony size and taxa on demographic fate

To demonstrate how colony-specific metrics predicted demographic fates, we evaluated how colony genus and size prior to the heatwave influenced the probability of colonies undergoing each demographic fate. We fit a multinomial logistic regression of colony fate (Growth, Stasis, Partial mortality, or Complete mortality) as a function of the interaction between pre-disturbance colony size and genus for all *Acropora* and *Pocillopora* colonies (n = 3,162), with growth specified as the reference outcome. Colony size was natural-log transformed prior to analysis. We evaluated whether the relationship between initial colony size and demographic fate differed between genera by comparing the full interaction model with an additive model containing only the main effects of colony size and genus using a likelihood-ratio test and Akaike’s information criterion (AIC). We retained the interaction model based on improved model fit. From the fitted model, we estimated the probability of each demographic fate and associated 95% confidence intervals across the observed range of colony sizes for each genus. Multinomial models were fit using the multinom() function in the nnet package (Venables & Ripley 2002), and predicted probabilities and confidence intervals were estimated using the emmeans package (Lenth 2026) in R.

#### Tracking multi-year colony fate trajectories

For a single 25 m^2^ reef plot, we explored change over a four-year period to demonstrate the application of our method for tracking multi-year colony fates (for *Pocillopora* only). In addition to the two-year period that spanned the heatwave event (2018-2019), this extended time series included annotations for 2017 (one year and eight months before the heatwave) and 2020 (one year and four months after the heatwave). This expanded analysis illustrates benthic dynamics during both a period without disturbance (2017-2018), as well as delayed impacts of the heatwave that occurred during a period after heat stress had fully subsided (2019-2020). We tracked the condition of each *Pocillopora* colony > 20 cm^2^ that was alive at the initial time point (n = 425) as ‘Live’, ‘Partially dead’, or ‘Completely dead’ through all four years, establishing a four-year trajectory (fate) for each colony. We then categorized each unique four-year fate, tracking the number of colonies that fell into each.

#### Changes in size structure

Finally, we quantified the size distributions for all live, partially dead, and completely dead *Pocillopora* colonies in each year to demonstrate changes in size structure of coral colonies over the extended study period.

All statistics and visualizations for this study were conducted in R (Version 4.2.3; R Core Team, 2023) and RStudio (Version 2023.12.1.402; Posit team, 2024) and utilized colors from the Manu New Zealand Bird Colour Palettes (Thomson, 2022). Some code was developed with the assistance of ChatGPT (version 4.0) to identify statistical approaches and their proper reporting format, as well as develop reproducible visualizations.

## RESULTS

### Tracking colony fates through a disturbance event

Our method enabled tracking the demographic fates of 2931 colonies of *Pocillopora* spp. (collectively, 67.6 m^2^ in live tissue surface area), and 231 colonies of *Acropora* spp. (5.2 m^2^ in live tissue surface area) across five ∼25 m^2^ reef plots through a heatwave-induced mortality event (Fig. 3A). 1186 colonies (40.4%) of *Pocillopora* and 163 colonies (70.6%) of *Acropora* underwent Complete mortality during this period. Together, Complete mortality accounted for a loss of 36.0 m² of living coral tissue (Fig. 3B). A further 627 colonies (21.4%) of *Pocillopora* and 23 colonies (10%) of *Acropora* underwent Partial mortality, accounting for an additional loss of 6.6 m^2^ and 0.2 m^2^ of live coral for each taxon, respectively. A substantial number of *Pocillopora* colonies, 915 (31%), survived and grew during this period, but this accounted for an increase of only 4.4 m^2^ of live coral tissue. Only 23 colonies (10%) of *Acropora* increased in size, contributing a gain of only 0.2 m^2^ of live coral. A further 203 colonies (7%) of *Pocillopora* and 16 colonies (7%) of *Acropora* exhibited Stasis during this one-year period.

**Figure 3.**
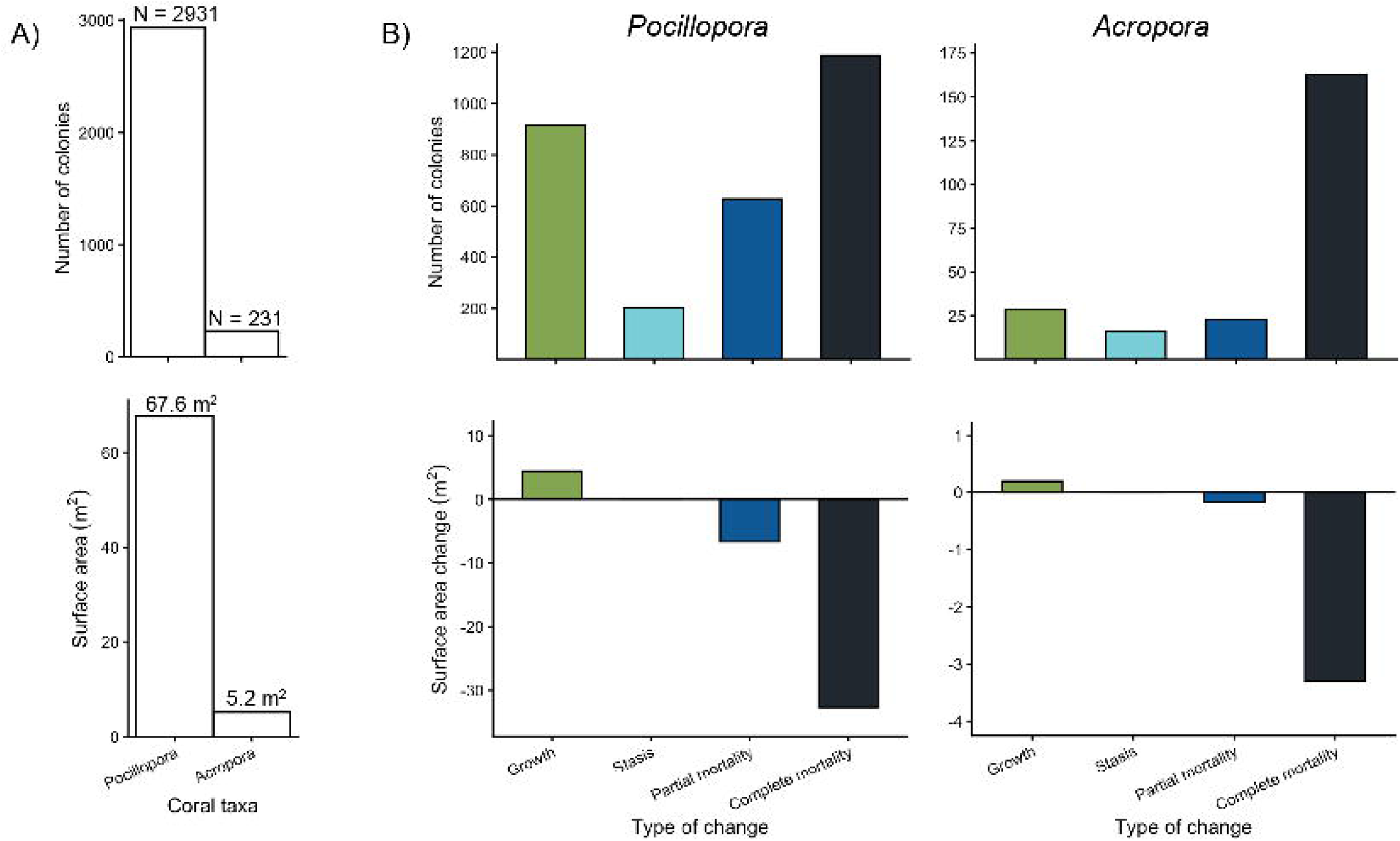
(A) Total number of colonies (top) and their collective 3-D surface area (bottom) for *Pocillopora* and *Acropora* across five replicate reef plots (each ∼25 m² in area) in 2018 prior to the marine heatwave. (B) The number of colonies from the two taxa that fell into each category of change (fate) from 2018 to 2019, and the change in surface area of live coral tissue associated with each change category. Colors associated with each fate category are included for consistency with subsequent figures.

### Influences of colony size and taxa on demographic fate

Our workflow revealed size- and genus-dependent patterns in demographic fates resulting from the heatwave (Fig. 4). The relationship between initial colony size and demographic fate differed between *Pocillopora* and *Acropora* (size × genus interaction: likelihood-ratio χ² = 26.00, *df* = 3, *P* < 0.001; ΔAIC = 20.00). For *Pocillopora*, the predicted probability of Growth declined from 0.39 (95% CI: 0.36–0.42) for colonies 50 cm² in surface area to 0.20 (0.17–0.23) for colonies 1,000 cm² in surface area, while the probabilities of Stasis, Partial mortality, and Complete mortality increased with colony size (Fig. 4A). In contrast, Complete mortality was strongly size-dependent in *Acropora*, declining from a predicted probability of 0.89 (0.81–0.97) for 50 cm² colonies to 0.28 (0.07–0.49) for 1,000 cm² colonies, while the probability of Stasis increased from <0.01 to 0.41 (0.06–0.77). Among colonies that survived the heatwave, changes in colony surface area ranged from substantial partial mortality to large increases in size, with most colonies experiencing more moderate changes (Fig. 4B). Complete mortality occurred across a broad range of initial colony sizes in both genera, with the greatest numbers of completely dead colonies occurring at small to intermediate sizes (Fig. 4C), reflecting the underlying size distribution of colonies as well as size-dependent mortality probabilities.

**Figure 4.**
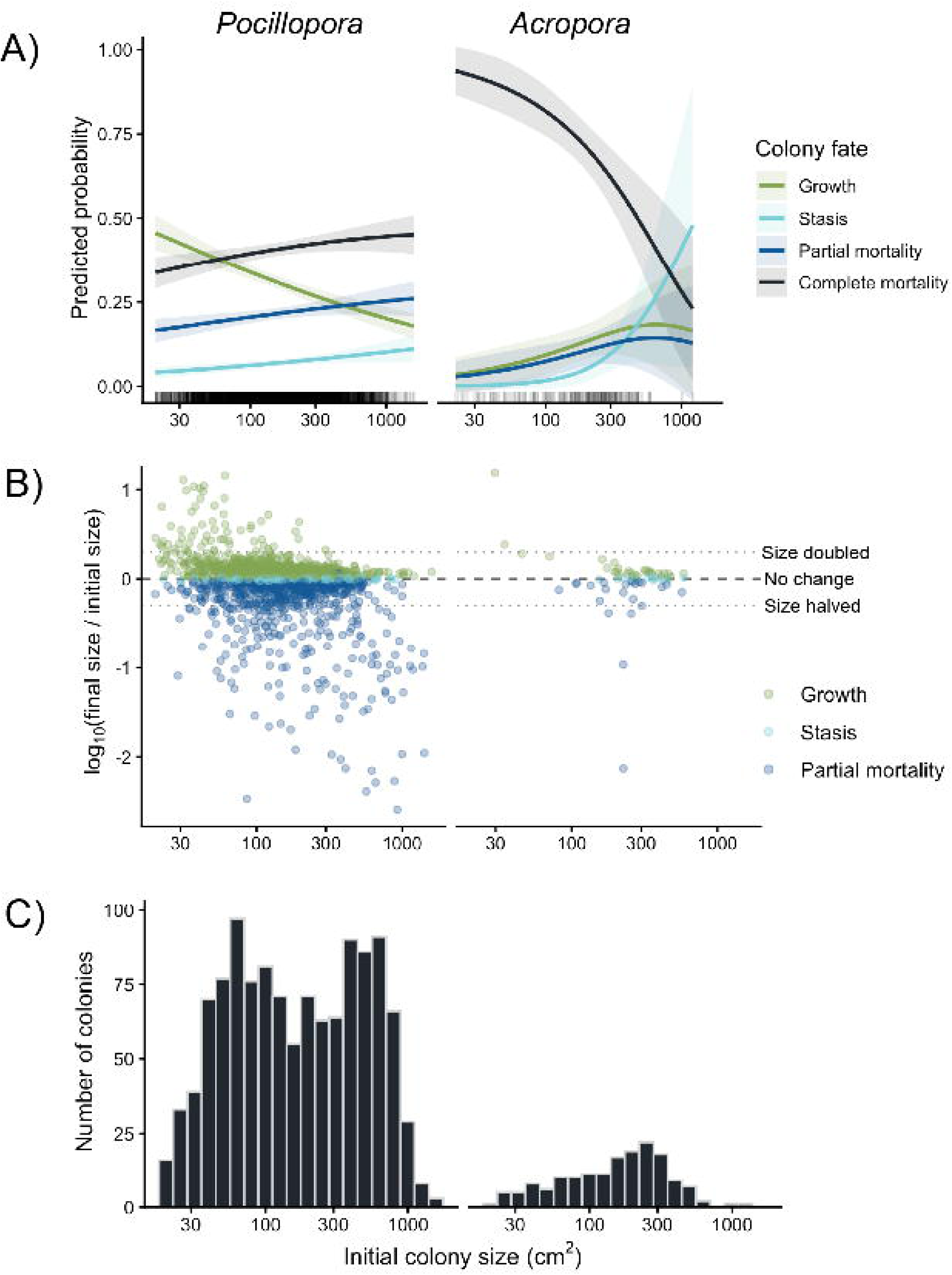
Colony-level demographic responses to the 2019 heatwave as a function of colony genus and pre-disturbance size. (A) Predicted probabilities of four demographic fates (Growth, Stasis, Partial mortality, and Complete mortality) from a multinomial logistic regression including the interaction between natural-log-transformed initial colony size and genus. Lines show model-predicted probabilities and shaded regions indicate 95% confidence intervals; rugs along the x-axis show the distribution of observed initial colony sizes for each genus. (B) Observed change in colony size among colonies that persisted through the heatwave, expressed as the log10 response ratio, log10(final size / initial size), and colored by demographic fate. A value of zero indicates no net change in colony size, while positive and negative values indicate net growth and loss, respectively. Horizontal dotted lines indicate a doubling [log10(2) = 0.301] or halving [log10(0.5) = −0.301] of colony size. (C) Distribution of initial colony sizes for colonies that experienced complete mortality. Initial colony size (x-axis) is displayed in cm² on a logarithmic scale in all panels for visualization purposes.

### Tracking multi-year colony fate trajectories

Within the single ∼25 m² plot used for our extended time series, the workflow revealed temporal changes in coral abundance and colony condition during both non-disturbance and disturbance periods. During the period without disturbance (2017-2018), the total surface area of live coral (*Pocillopora* + *Acropora*) increased by 0.7 m^2^ (5%) and dead coral increased by 0.6 m^2^ (Fig. S1). During the heatwave-induced mortality event (2018-2019), live coral decreased by 6.5 m^2^ (45%), resulting in an eightfold increase of standing dead coral cover within the plot. Notably, live coral continued to decrease in the year following the heatwave, declining by a further 1.4 m^2^ (18%) and increasing dead coral by the same amount (Fig. S1).

Analysis of changes in coral colony condition over four years revealed a variety of multi-year fate trajectories (Fig. 5). For the 425 live *Pocillopora* colonies present in the first time point, we tracked 10 unique fate trajectories through the three subsequent time points (Fig. 5, Table 1). Of these, 193 (45.4%) survived either partially or completely until 2018, then suffered complete mortality during the heatwave. In contrast, 123 colonies (28.9%) remained at least partially alive through the final time point, representing three of the 10 unique fate trajectories (Table 1). This group of colonies – i.e., those that were either completely or partially alive by the final time point – represented 3 of the 10 unique fate categories (Table 1). 94 of these colonies (22.1%) showed no signs of mortality through all four years and either grew or exhibited stasis throughout the study period. 32 colonies (7.5%) underwent ambient mortality between the first and second years – i.e., mortality that was not induced by the heatwave. Finally, 77 colonies (18.1%) died completely during the year after the heatwave (2019-2020; Table 1).

**Figure 5.**
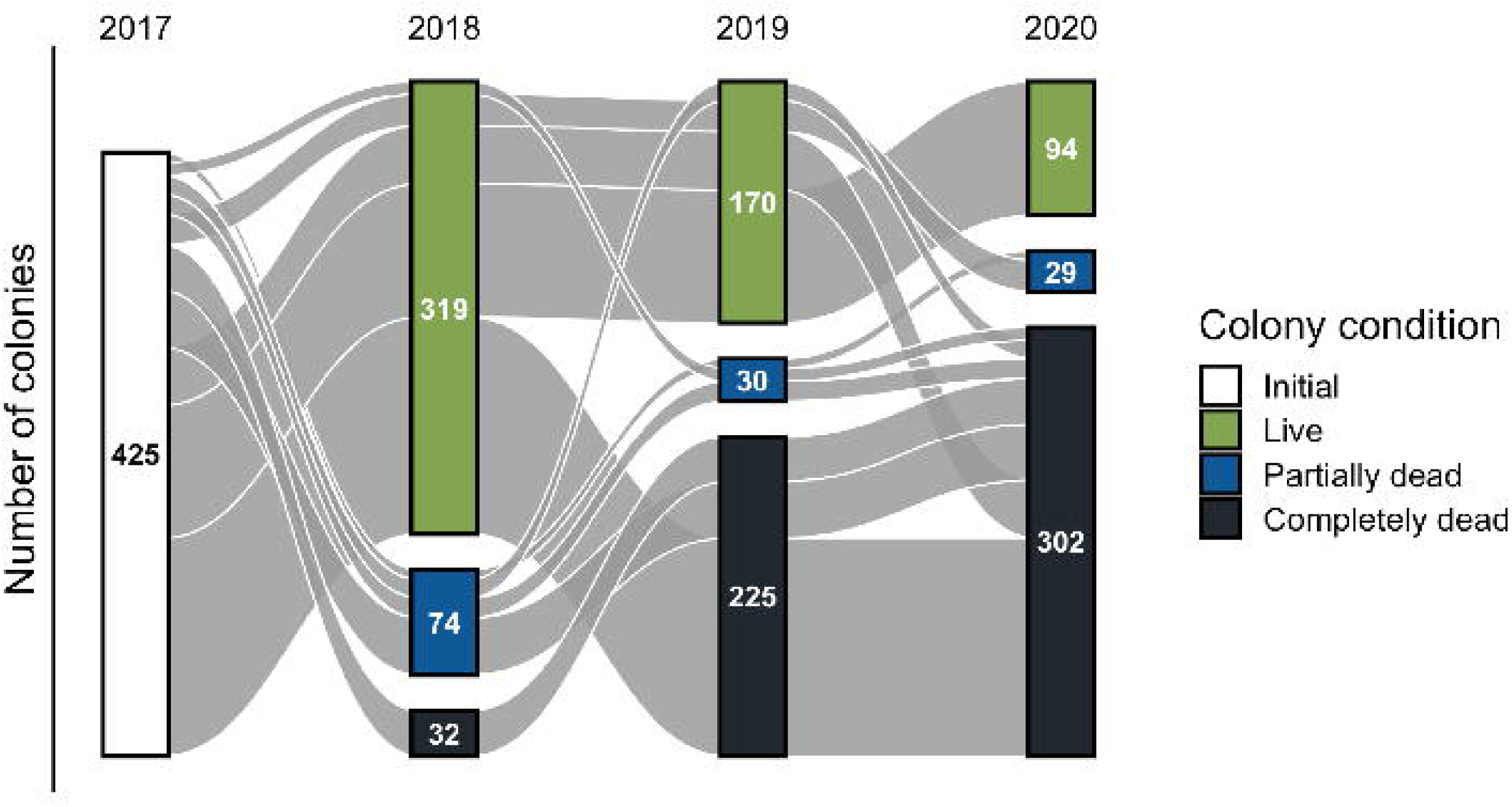
Sankey plot displaying changes in colony condition through time (i.e., multi-year fates) of all *Pocillopora* colonies that were alive in 2017 within a single 25 m² reef plot over four years (n = 425 colonies). Nodes (vertical bars) represent the number of colonies in each year that were completely alive (including colonies that either grew or underwent stasis during the previous year), partially dead, or completely dead. Edges (horizontal flows) show transitions of colonies among these categories through time. Edges (n = 10) group together all colonies that followed the same four-year fate (see Table 1 for exact counts). Edges (fates) that contained fewer than five colonies were omitted for visualization purposes.

**Table 1.** Four-year fates of coral colonies that were initially alive in 2017 within a single ∼25 m² reef plot (n = 425). Each row represents a unique four-year fate trajectory and indicates both the number and percentage of colonies that underwent each fate. Fates with fewer than five colonies were omitted to align with Figure 5.

| 2017 | 2018 | 2019 | 2020 | Colonies | Percent (%) |
| --- | --- | --- | --- | --- | --- |
| Live | Live | Completely dead | Completely dead | 153 | 36.0 |
| Live | Live | Live | Live | 94 | 22.1 |
| Live | Live | Live | Completely dead | 41 | 9.6 |
| Live | Partially dead | Completely dead | Completely dead | 40 | 9.4 |
| Live | Completely dead | Completely dead | Completely dead | 32 | 7.5 |
| Live | Live | Live | Partially dead | 22 | 5.2 |
| Live | Partially dead | Partially dead | Completely dead | 14 | 3.3 |
| Live | Partially dead | Live | Completely dead | 13 | 3.1 |
| Live | Live | Partially dead | Completely dead | 9 | 2.1 |
| Live | Partially dead | Partially dead | Partially dead | 7 | 1.6 |

### Changes in size structure

The size structure of live Pocillopora colonies (>20 cm²) changed substantially over the four-year study period (Fig. 6). While colony abundance declined across size classes, the majority of living colonies that transitioned between size classes in successive time points did so by increasing rather than decreasing in size. Standing dead colonies also accumulated within the reef plot through time.

**Figure 6.**
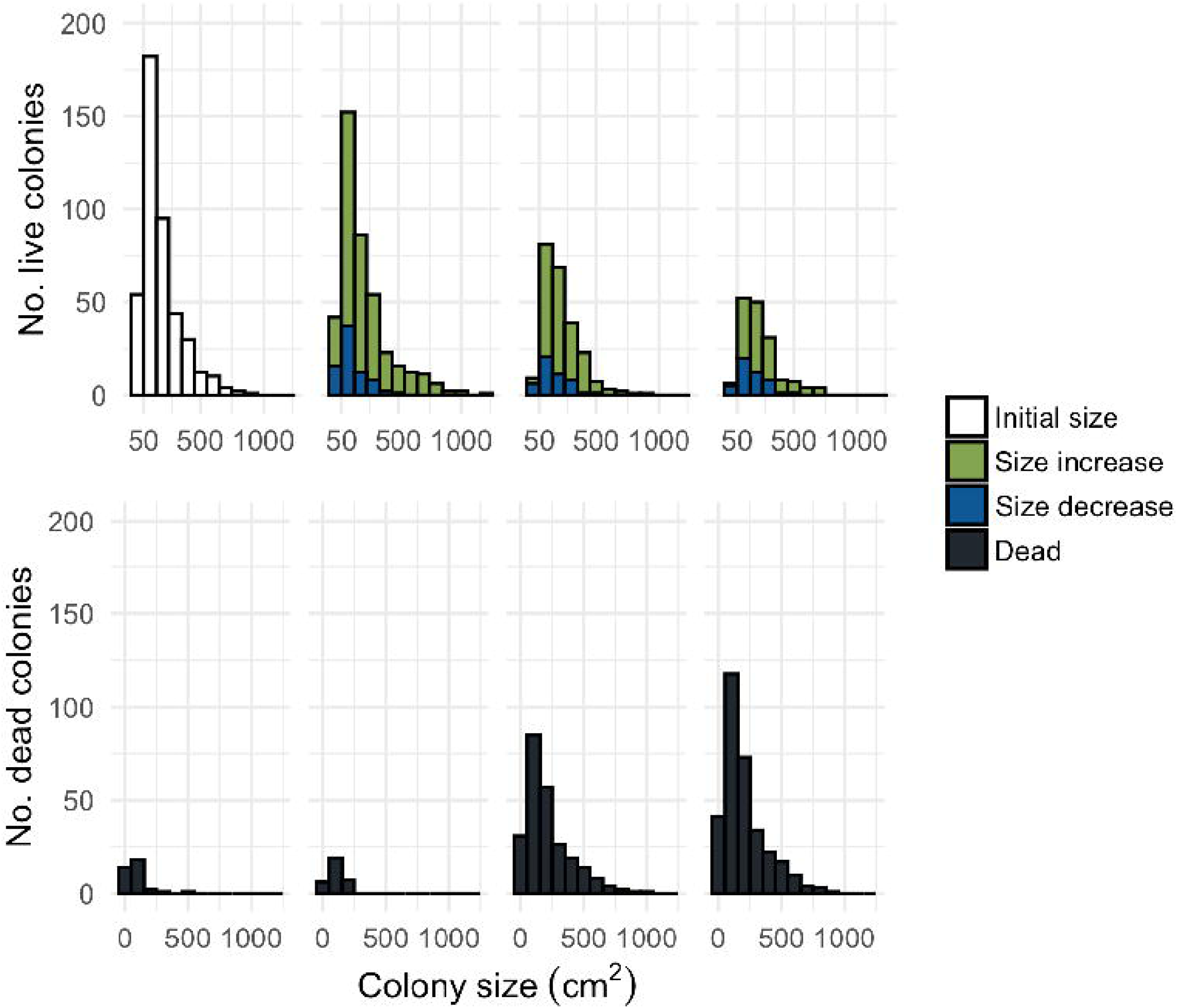
Size distributions of live (top) and dead (bottom) Pocillopora colonies within a single 25 m² reef plot over four years. In the top row, white bars indicate the size distribution of the initial cohort of live colonies in 2017, and green and blue bars in subsequent years indicate live colonies that either grew or shrank (i.e., died partially) since the previous year and thereby entered a new size bin. Near-black bars indicate dead colonies of various sizes in each year.

## DISCUSSION

The ability to link demographic processes that occur at the scale of individuals with the broader spatial areas they inhabit is an important step toward effectively quantifying demographic changes of space-holding organisms across a wide range of ecosystems. We present a workflow that combines underwater photogrammetry and ML-assisted image analysis to track changes in reef-building corals, and potentially clonal and non-clonal sessile organisms in other ecosystems. We demonstrate a subset of possible applications of this workflow for tracking annual demographic changes in hundreds to thousands of colonies across spatially replicated reef surveys, both in the absence and presence of disturbance-driven declines. By preserving individual identities through repeated surveys, our approach provides a means to integrate high-resolution demographic information with spatially extensive mapping of coral reefs and other benthic environments.

ML is increasingly implemented to automate the extraction of data from remote sensing imagery on coral reefs (C. Burns et al., 2022; Kopecky, Pavoni, et al., 2023; Sauder et al., 2024; Schürholz & Chennu, 2023; Stupariu et al., 2022). We build on these previous efforts by integrating image-based mapping with temporal matching of individual colonies, allowing repeated surveys to track demographic cahnges at the colony level. Compared to simply estimating net loss of coral cover following a disturbance, this approach provides a more nuanced assessment of disturbance impacts by showing how colony-level characteristics (pre-disturbance size, taxa) influence the likelihood of multiple demographic outcomes. We therefore gain a more informed understanding of how large numbers of corals are likely to vary in their responses to disturbance and identify characteristics associated with greater or lower vulnerability. Similar approaches are being applied in other systems to link demographic processes in slow-growing, non-mobile organisms such as trees from individuals to landscapes (Battison et al., 2024; Marconi et al., 2021; Weinstein et al., 2024). Technological tools and workflows such as these could help ecosystem managers identify groups of organisms that are particularly vulnerable to disturbance and inform the allocation of limited resources for intervention efforts. Both the matching algorithm in TagLab that enables these automated analyses of coral growth and mortality, as well as and the software as a whole, are publicly available and well documented for wide user accessibility.

The workflow we present here also addresses an enduring challenge in the study of coral populations—effectively quantifying partial colony mortality (Jackson & Hughes, 1985). This intermediate condition between mortality and survival makes accurately censusing coral populations difficult and conflates demographic mechanisms underlying population change, as corals can enter new size classes either by growing or by shrinking (i.e., partially dying). This challenge is not unique to corals, as other organisms exhibit clonal growth patterns and can similarly undergo partial mortality. For example, seagrasses and terrestrial grasses expand through rhizomatic growth and can lose parts of their biomass to various causes while retaining a live portion (Bell & Tomlinson, 1980; Marbà & Duarte, 1998). Our technique could be similarly useful for demographic studies and monitoring in these other systems by incorporating partial mortality as a metric to better understand temporal changes and disturbance impacts in clonal organisms.

There is mounting evidence surrounding the powerful influences of dead organisms on living ones (Kopecky et al., 2026; Saldaña et al., 2023); therefore, classifying both live and dead organisms will become increasingly important for understanding ecological impacts of disturbance. This is especially true in coral reef ecosystems (Kenyon et al., 2023; Kopecky et al., 2024, 2025; Kopecky, Stier, et al., 2023; Norström et al., 2007), where disturbances like marine heatwaves and outbreaks of coral predators that generate standing stocks of dead skeletons are becoming more prevalent (Byrne et al., 2024; Oliver et al., 2018; Pratchett et al., 2017). Our workflow enables automatic classification and measurement of both live and dead coral colonies, providing a means to make more complete assessments of changing reef health (Kopecky et al., 2025; Kopecky, Pavoni, et al., 2023).

While our workflow offers considerable advances for coral reef mapping and monitoring, we acknowledge some limitations remain. First, to automatically identify the same colonies (or other objects) across multiple time points, maps must be co-registered in space with a common, fixed reference system. Though this is straightforward in terrestrial or shallow water systems where GPS can be used, deeper water habitats (where GPS is ineffective) require installation of permanent reference networks (see Nocerino et al., 2020). While this requirement places constraints on the sizes of reef tracts that can be feasibly surveyed, more advances are underway to reduce this barrier (Nocerino & Menna, 2023). Second, because creating reliable automatic classification algorithms requires sufficient training data, this method performs well on common taxa but less so on rarer taxa. Therefore, holistic censuses of benthic (or sessile) communities would require some additional manual annotation and user effort to create training datasets describing both common and rare taxa. Still, the semi-automatic segmentation tools used here can reduce the manual effort required for annotation (Pavoni et al., 2022). Lastly, accurately delineating the boundaries between overlapping coral colonies of the same class (i.e., live overlapping live or dead overlapping dead) remains a challenge in the automatic classification process. This can result in overestimation of colony sizes and issues with matching colonies between time points. This occurred for only a small proportion (< 10%) of the automatically classified colonies we followed, and these issues were easily resolved through semi-automated editing of colony borders.

Understanding the consequences of global change is among the most important endeavors in modern ecology. Our workflow provides an avenue for integrating individual-level demographic information with repeated image-based monitoring of coral reefs, as well as other ecosystems characterized by sessile organisms where photogrammetry is possible. By linking demographic changes in individual organisms with spatially extensive, repeated observations, approaches such as these can contribute to a more comprehensive understanding of how ecosystems respond to rapidly changing environmental conditions.

## ACKNOWLEDGMENTS

We thank Hillary Krumbholz for assistance with Information Management and field support, Jordan P. Gallagher, Dana T. Cook, and Randi N. Honeycutt for their invaluable technical assistance in the field, and the staff of the University of California Gump Research Station and UC Dive & Boat Safety Enterprise for logistic support. Field research was completed under permits issued by the Territorial Government of French Polynesia (Délégation à la Recherche) and the Haut-commissariat de la République en Polynésie Française (DTRT) (Protocole d’Accueil 2006-2026); we thank them for their continued support. With respect to the place name spelling of Moorea in this paper, we followed the Raapoto transcription system for Reo Mā’ohi (the traditional language of the Society Islands) that is adhered to by a large segment of the Tahitian community, but we also recognize other community members follow the Te Fare Vanā’a transcription system where the island name is spelled with an ’eta (Mo’orea). This work was supported by the U.S. National Science Foundation (OCE 2224354 and earlier awards for the Moorea Coral Reef LTER); the Gordon and Betty Moore Foundation; the Italian Ministry of University and Research (PNRA18-00263-B2); and ETH Zurich.

## DATA ACCESSIBILITY

All data and code scripts used for analyses and visualizations presented in this study can be found at the following GitHub repository: https://github.com/kkopecky711/Coral-colony-fates.git. Upon acceptance of this manuscript, data will be permanently archived with the

Environmental Data Initiative (EDI), and both the data and coded workflows will be permanently and publicly archived both in Zenodo and GitHub.

## DECLARATION ON THE USE OF AI

Some code was developed with the assistance of ChatGPT (version 4.0) to identify statistical approaches and their proper reporting format, as well as develop reproducible visualizations.

## AUTHOR CONTRIBUTIONS

KLK, RJS and SJH conceived the ideas; KLK, GP, MC, EN, FM, AJB, RJS and SJH designed the methodology and conducted the investigation; KLK, GP, MC, EN and FM curated and analyzed the data; KLK, GP and MC developed the software; KLK created the visualizations; GP, MC, EN, FM, AJB, RJS and SJH provided resources and acquired funding; AJB, RJS and SJH provided supervision; KLK led the writing of the manuscript. All authors contributed to project administration and validation, contributed critically to the drafts, and gave final approval for publication.

## STATEMENT ON INCLUSION

Our study brings together authors from several different countries, but due to a lack of researchers working with the specific tools and methodologies we describe here, our study does not include scientists based in the country where the study was carried out. We do, however, discuss our workflow with local stakeholders whenever possible and appropriate to seek feedback, engage in knowledge exchanges, and make our efforts and methods both known and available to local communities in the locale of our research for uses they see fit.

## AUTHORS’ DECLARATION

All authors have seen and approved the submitted version of the manuscript and have substantially contributed to the work. All persons entitled to co-authorship have been included as authors. The authors confirm that this manuscript has been submitted solely to Remote Sensing in Ecology and Conservation and has not been published elsewhere, in whole or in part, nor is it currently in press or under consideration for publication by another journal.

## CONFLICT OF INTEREST

All authors declare no conflict of interest.

## Notes

https://github.com/kkopecky711/Moorea-hysteresis-and-resilience-experiments

### Competing Interest Statement

The authors have declared no competing interest.

