## Supporting information for "Tracking demographic changes in individual coral colonies through repeated photogrammetric surveys"

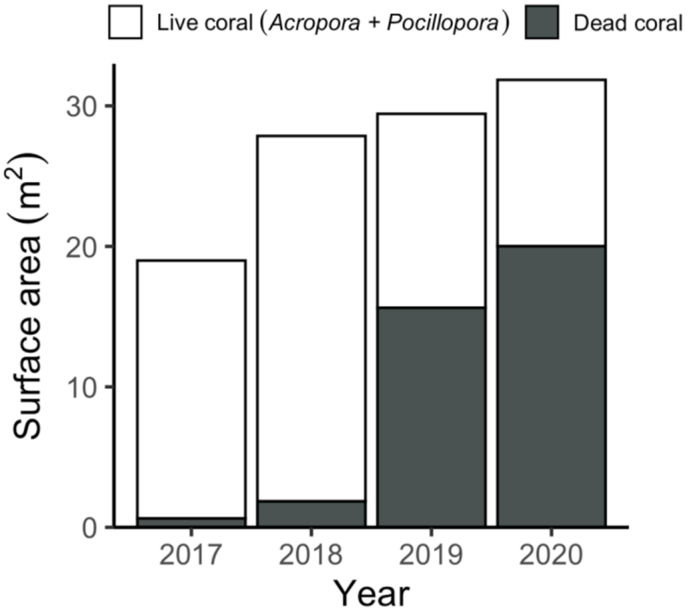


**Figure S1.** Total 3-D surface area of live coral (*Acropora* and *Pocillopora* combined) and dead coral for a single (~25 $m^2$) reef plot from 2017-2020.

**Table S1.** Multinomial logistic regression of coral colony demographic fate as a function of initial colony size and genus. Growth was set as the reference demographic fate and *Pocillopora* as the reference genus. Initial colony size was natural-log transformed. Estimates therefore represent changes in the log odds of each demographic fate relative to growth. *P*-values are based on Wald z-tests.

| Fate | Term | Estimate | SE | z | p |
|---|---|---|---|---|---|
| <chr> | <chr> | <dbl> | <dbl> | <dbl> | <dbl> |
| Stasis | (Intercept) | -3.75 | 0.457 | -8.20 | 2.50e-16 |
| Stasis | log(area_2018) | 0.443 | 0.0872 | 5.09 | 3.66e- 7 |
| Stasis | classAcropora | -3.18 | 3.33 | -0.956 | 3.39e- 1 |
| Stasis | log(area_2018):classAcropora | 0.685 | 0.584 | 1.17 | 2.41e- 1 |
| Partial mortality | (Intercept) | -1.97 | 0.297 | -6.63 | 3.35e-11 |
| Partial mortality | log(area_2018) | 0.319 | 0.0583 | 5.47 | 4.51e- 8 |
| Partial mortality | classAcropora | 1.79 | 2.31 | 0.775 | 4.38e- 1 |
| Partial mortality | log(area_2018):classAcropora | -0.328 | 0.425 | -0.772 | 4.40e- 1 |
| Complete mortality | (Intercept) | -1.14 | 0.250 | -4.57 | 4.98e- 6 |
| Complete mortality | log(area_2018) | 0.281 | 0.0496 | 5.67 | 1.39e- 8 |
| Complete mortality | classAcropora | 6.72 | 1.66 | 4.06 | 4.95e- 5 |
| Complete mortality | log(area_2018):classAcropora | -1.02 | 0.307 | -3.32 | 9.07e- 4 |
